# Development of an ultrafast pulsed ponderomotive phase plate for cryo-electron tomography

**DOI:** 10.1101/2024.03.20.585981

**Authors:** Daniel X. Du, Adam C. Bartnik, Cameron J. R. Duncan, Usama Choudhry, Tanya Tabachnik, Chaim Sallah, Ebrahim Najafi, Ding-Shyue Yang, Jared M. Maxson, Anthony W. P. Fitzpatrick

**Affiliations:** Mortimer B. Zuckerman Mind Brain Behavior Institute, Columbia University, New York, NY 10027, USA; Department of Biochemistry and Molecular Biophysics, Columbia University, New York, NY 10032, USA; Taub Institute for Research on Alzheimer’s Disease and the Aging Brain, Columbia University Irving Medical Center, 630 West 168th Street, New York, NY 10032, USA; Cornell Laboratory for Accelerator-Based Sciences and Education, Cornell University, Ithaca, New York, NY 14853, USA; Università degli Studi di Milano-Bicocca, Milan, Italy; Department of Mechanical Engineering, University of California, Santa Barbara, CA 93106, USA; The Chemours Company, Wilmington, DE 19899, USA; Department of Chemistry, University of Houston, Houston, TX 77204, USA

## Abstract

Cryo-electron tomography (cryo-ET) is a powerful modality for resolving cellular structures in their native state. While single-particle cryo-electron microscopy (cryo-EM) excels in determining protein structures purified from recombinant or endogenous sources, cryo-ET suffers from low contrast in crowded cellular milieux. A novel experimental approach to enhance contrast in cryo-ET is to manipulate the phase of scattered pulsed electrons using ultrafast pulsed photons. Here, we outline the experimental design of a proof-of-concept electron microscope and demonstrate synchronization between electron packets and laser pulses. Further, we show ultrabright photoemission of electrons from an alloy field emission tip using femtosecond ultraviolet pulses. These experiments pave the way towards exploring the utility of the ponderomotive effect using pulsed radiation to increase phase contrast in cryo-ET of subcellular protein complexes *in situ*, thus advancing the field of cell biology.

## Introduction

Amino acid sidechains were first resolved by single particle cryo-EM imaging of viruses on photographic film in 2008^1,2^. By 2013, the advent of direct electron detectors ushered in an era of “near-atomic” cryo-EM (better than 4 Å resolution) using single particle analysis^3,4^. While it has been demonstrated that cryo-ET can attain similar resolutions *via* subtomogram averaging, cellular crowding and the presence of low individual protein copy numbers present a challenge to the detection, identification, and determination of structures within cells^5–7^. One experimental approach to enhancing contrast in cryo-EM images, akin to a Zernike phase plate, is to alter the phase of a post-specimen electron beam through elastic interactions with photons^8^. The use of a ponderomotive field potential, created by focused laser beams to modulate the phase of a transmitted electron beam, has advantages over post-specimen material phase plates which attenuate the transfer of high-spatial frequencies and suffer from charging and damage^9–15^.

Recently, this concept which requires high peak intensities has been realized in two ways. The first approach employs a Fabry-Perot resonator (Fig. 1a), while the second utilizes a ring micro-resonator (Fig. 1b)^14,16^. Images of biological samples show contrast enhancements comparable to those achieved with material phase plates^17–19^. Improvements to resonator-based laser phase plates center on; (i) attempting to correct spherical and chromatic aberrations arising from scattering^20^, and (ii) minimizing obtrusive and expensive physical alterations to the column of the transmission electron microscope^21^. Significantly, pulsed laser beams generate sufficiently high peak intensities to interact with a pulsed electron beam^22,23^. Pulsed laser beams, which enter the microscope externally through optical windows (Fig. 1c), offer a practical alternative to resonators. Here, our stroboscopic experiments demonstrate highly reproducible spatial and temporal overlap of pulsed photons with photoelectron packets, necessary for future experiments on pulsed laser-electron interactions. We also discuss the capabilities of high-frequency pulsed lasers and next-generation photocathodes, with an emphasis on matching the brightness of modern-day field-emission-guns (FEGs).

**Fig. 1.**
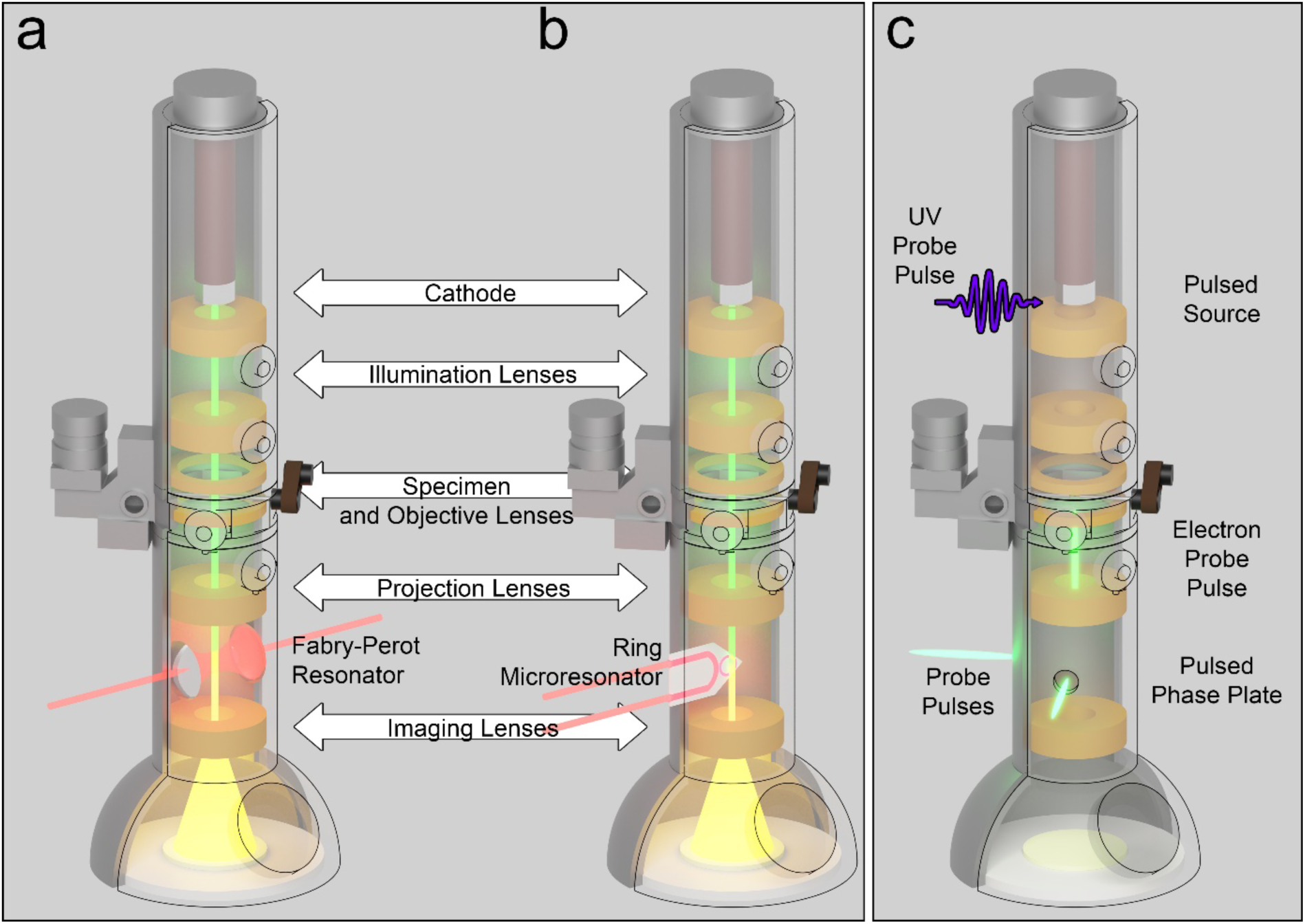
Schematics of the various laser-based phase plates. **a and b**, TEM schematics that demonstrate resonator phase-plates alongside potential locations for accessing the back focal plane using a (a) Fabry-Perot resonator^16^ or (b) a ring microresonator^14^. **c,** A demonstration of the pulsed laser phase plate, where pulses of electrons and photons overlap at a pre-determined location in the TEM to generate a phase shift in the electrons not scattered by the specimen^22^.

### Pulsed laser phase plate room layout

The layout of the pulsed laser phase experimental lab accommodates a scanning electron microscope (SEM), an optical table, and a transmission electron microscope (TEM)^24^. We have arranged each of these experimental components (Fig. 2) so that each electron microscope is accessible by the laser. The laser is oriented such that proof-of-principle experiments can be performed by interfacing an optical pump-probe scheme with the SEM. In particular, the generously sized low-vacuum (10^-4^ Pa) SEM chamber easily allows the placement of optical components and is adaptable for modifications. The laser (Fig. 2a, b) is an NKT Photonics Aeropulse FS20-050 fiber laser (400 fs pulse width) with a fundamental wavelength of 1030 nm that is frequency doubled to 515 nm. As is the case with many ultrafast and interferometry experiments, a beamsplitter is used to preserve the timing between pump and probe pulses and sidestep the issue of laser emission timing jitter^25^. Here, a 90:10 T:R plate beamsplitter sends 90% of the power through the pump line for specimen photoexcitation and 10% of the power to the probe line. In both lines, a waveplate and glan-calcite polarizer combination is used to modulate the power reaching the SEM. The probe line uses a Keplerian telescope and a Type 1 BBO placed at the focal point of the first lens to frequency double the 515 nm pulses into 257 nm pulses. The UV pulses are then sent upwards towards the optical port on the side of the SEM column that leads directly to the ZrO coated tungsten cathode, inducing photoemission of ultrashort electron pulses that travel down the column (Fig. 2c). The beam radii (ω) are focused down to a 7 µm × 14 µm spot using a 150 mm lens with an effective focal length of 137 mm. The beam waist is maintained at the tip to ensure optimal photoemission (Supplementary Fig. 1). The pump pulses are passed through a Galilean telescope to expand the beam for later focusing. A retroreflector on a motorized delay stage is used to control the temporal delay between the pump and probe pulses at the specimen. The pump passes into the SEM through a custom optical port with a Vis-AR coated 1.5” window secured by silicone glue and two concentric O-rings.

**Fig. 2.**
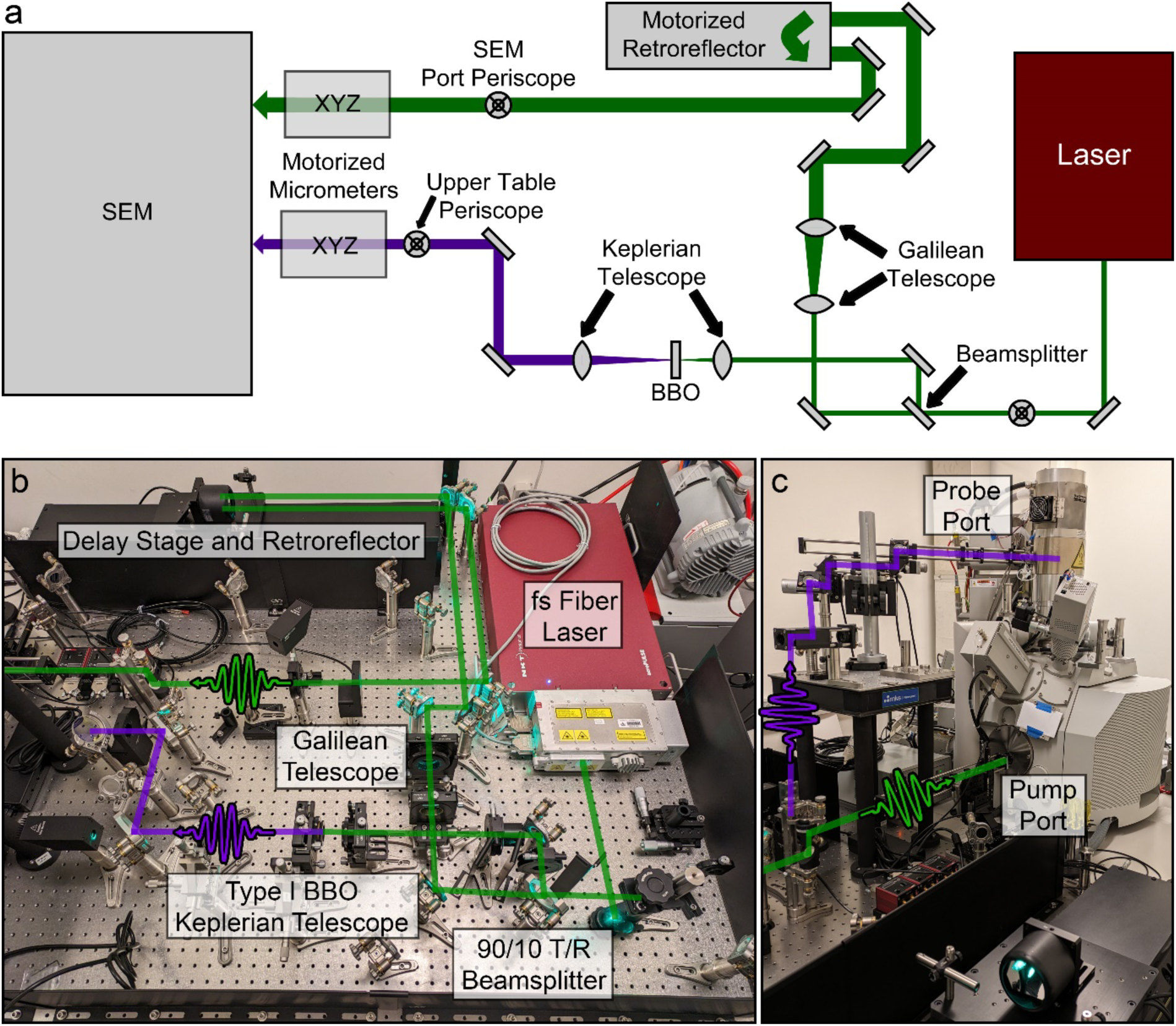
Overview of the laser table and interfaces with the SEM. **a**, A simplified schematic of the laser table with relevant beam modification hardware. The 515 nm laser signal (green line) originates from the SHG module on the fiber laser before being split in a 9:1 ratio. 10% of the power is directed into a Type 1 BBO, converting it into a 257 nm UV pulse (purple line). 90% of the power is directed into a retroreflector mounted on a motorized delay stage before being sent into the SEM. **b,** Implemented top-down view of the laser table, with various critical components and laser paths labeled. **c,** The two critical SEM ports are labeled, with dashed lines demonstrating how the green and UV laser pulses enter the system. The UV pulse is directed onto the SEM cathode, generating pulses of photoelectrons down the column. The green pulse is directed into an optical port that leads to internal periscopes that eventually reach the specimen.

### Photoexcitation of p-type silicon

Here, we demonstrate a scanning ultrafast electron microscopy (SUEM) experiment in a geometry that is easily transferrable to future experiments involving the ponderomotive effect (Fig. 3)^22^. A periscope with an adjustable lens mount is used to roughly align the laser into the electron beam field of view (Fig. 3a). Secondary and backscattered electrons are collected by an Everhart-Thornley Detector (ETD) (Fig. 3b). Critically, this experiment operates stroboscopically, with the signal in a single trial timepoint collected over millions of photoexcitation events^26^. The 2 MHz pump laser was set to a size of 11 µm × 16 µm, resulting in a 11 µm × 27 µm beam at the 30° tilted degenerate p-type boron doped (>10^19^ cm^-3^) silicon. The pulse energy *per* shot was 0.57 nJ, with a fluence of 122 µJ/cm^2^. Images were cyclically obtained from a reference timepoint well before time zero (-446 ps). Each cycle consisted of a forward and backward progression on the delay stage and a total of 20 cycles *per* trial position. This back-and-forth cycle served to average out the slowly decaying photoemission current and irreversible effects on the specimen. A different spot on the specimen was sampled for each trial. At each timepoint, images were formed at 500 ns dwell time *per* pixel across a 512 × 768 field of view and averaged 128 times before progressing to the next timepoint. The rapid acquisition of each image also minimized the accumulation of damage between timepoints.

**Fig. 3.**
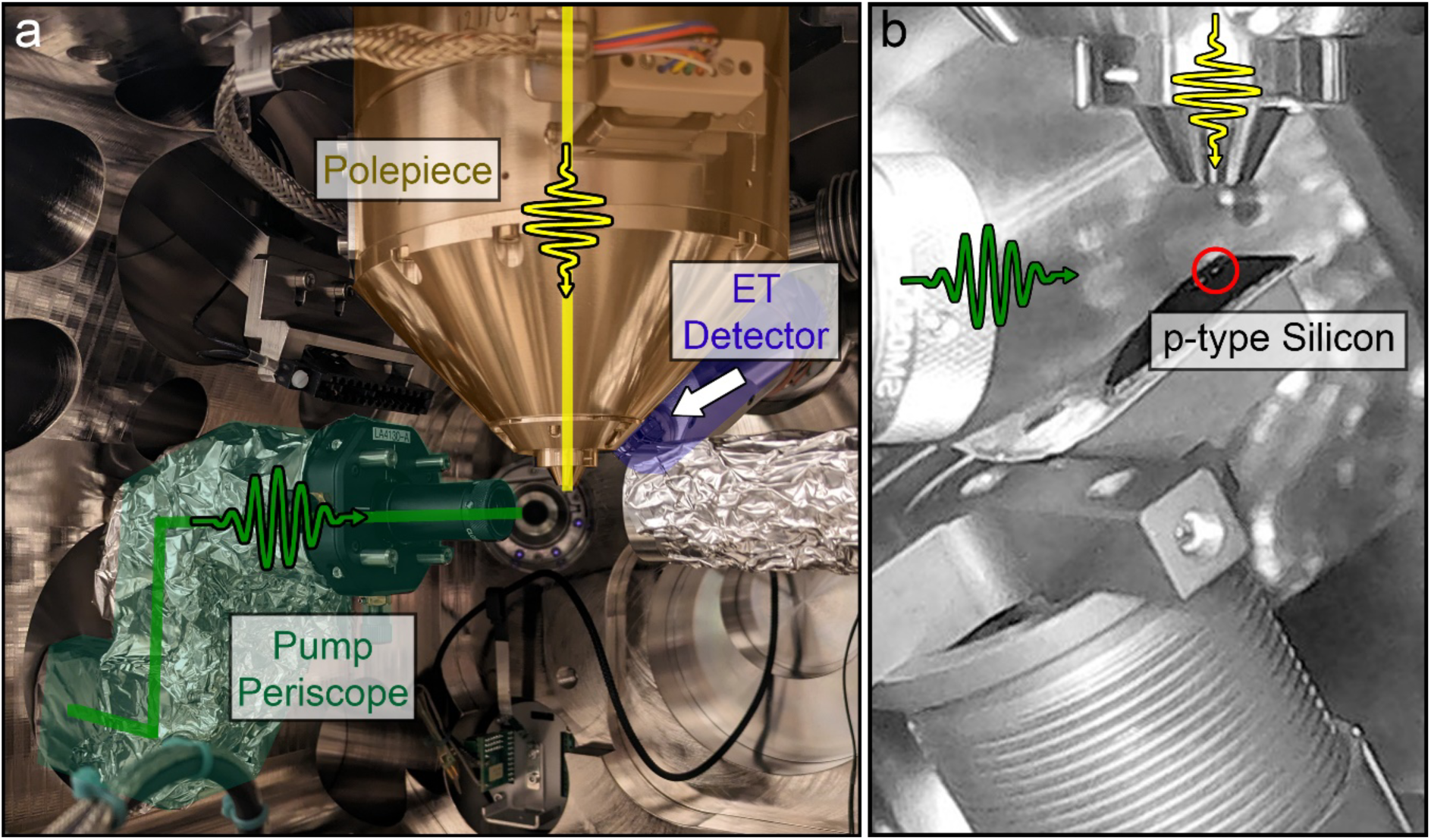
Internal experimental configuration of the SUEM. **a**, An image of the internal optics and polepiece. The laser (green dashed line) and electron (yellow dashed line) pulses intersect at the specimen position. The ETD is placed just behind the polepiece. **b,** An experiment with degenerate p-type silicon held in place on the standard SEM cross holder by a large piece of copper tape strongly adhered to both surfaces to prevent charging. The specimen is tilted 30° to allow both the laser and electrons to interact with the specimen. The interaction region is highlighted with a red circle. The pump pulse (green) photoexcites a response from the specimen. The electron pulse (yellow) probes a point along the response.

To adjust for variations in specimen position, each difference image was normalized to the total difference image intensity at an arbitrary time with a strong response. A reference image has been included in Supplementary Figure 2a. In this manner, many trials could be stitched together to form a movie containing 26 frames with a total duration of 3 ns. This averaging also minimizes the influence of variations in specimen surface condition. After image integration and normalization, the difference images showed several distinct signals (Fig. 4). The first signal is a slight rise in counts (secondary electron surplus) surrounding the laser spot beginning at time zero and continuing to rise until 87 ps, where a strong bright annulus that clearly appears in an area larger than the laser spot size. This bright signal then rapidly expands across 400 ps before reaching an equilibrium size that persists to the end of the available space on the delay stage, around 2.5 ns after time zero. The second signal is a dark region within the strong bright pattern that matches the size of the laser spot and first appears at 154 ps, also persisting until the end of the scan. It is believed that this dark area is a consequence of residual laser damage on the specimen as explored in the analysis section.

**Fig. 4.**
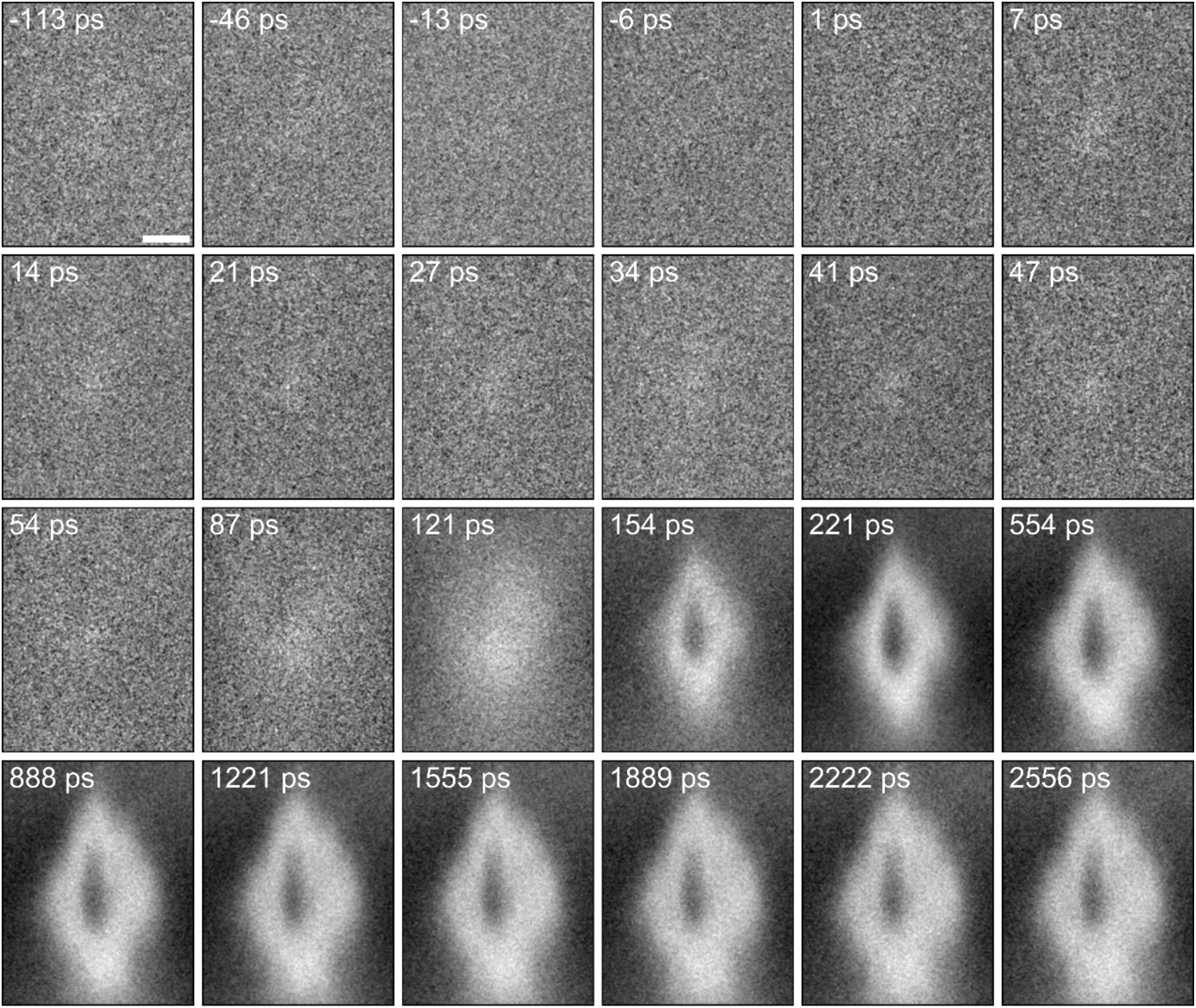
Difference images demonstrating the signal arising from photoexcitation. A series of difference images relative to a reference time point well before time zero. The initial rise is first observed as a general increase in counts in a large area surrounding the laser spot. After 50 picoseconds, this brightness then evolves into a well-defined spot that persists for longer than 2.5 ns. The scalebar provided in the reference frame is 25 microns. The strong signal is characterized by a bright oval larger than the laser spot size surrounding a dark region approximately the size of the laser. It is believed that the darker region is caused by damage to the surface of the specimen. The bright region is a response from the disruption in the equilibrium Fermi distribution.

The sum of each difference image in the region of interest was plotted against delay to clearly visualize the progression of signal strength (Fig. 5). Given that the resulting data shows no clear decay in the timepoints available to the delay stage, a sigmoid of the form

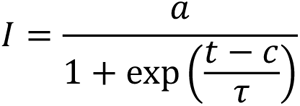

was chosen as an empirical fit to estimate the rise time of the signal, with *a* = 1.42 ± 0.05, *c* = 143 ± 14 ps, and τ = 49 ± 9 ps. At its peak, the difference represents an increase of 38% in counts within the image.

The lack of any decay in the available scan range is expected based on charge-carrier ultrafast studies reporting long-lived decays that occur between 100 and 200 ns^27,28^. More importantly, the rise time of the signal itself occurs over 49 ps. A result reported by Mohammad *et. al* showed the rise time of these signals increased linearly to pump pulse energy. This line indicates that the pulse energy of 600 nJ used in our experiment should have a rise time of 200 ps^29^, though such times are also affected by the photoelectron pulses that reach the specimen. The number of electrons *per* pulse influences the pulse duration of the electron beam based on the deleterious charge-charge effects on electron pulse duration. This will further affect the rise time of the signal. Previous measurements show that the high UV beam pulse energy of 7.5 nJ used in this experiment suggests additional rise times of at least 122 ps, though this can drastically change based on tip and electron microscope condition^30^.

Indeed, the dark contrast feature that appears at 154 ps and is the approximate size of the laser spot is inconsistent with another paper published by Najafi *et. al.*, where the signal at fluences from 160 µJ/cm^2^ to 1280 µJ/cm^2^ featured no sudden dark contrast feature, though the rise time was consistent with our results^31^. In another paper published by Liao *et. al.* and involving experiments at 20 µJ/cm^2^ to 67 µJ/cm^2^ pump pulses, a similar peak signal time was observed at 50 ps^32^. Additionally, the center of the signal showed a significant decrease in counts, similar to our result. There is also some confusion in the literature over whether p-type silicon produces an overall dark contrast feature, as observed in Li *et. al.* or a bright contrast feature^31,33^. Ultimately, there is more to be explored to understand the fluctuations in secondary electron emission^34^, though the data does clearly show ultrafast time-resolved dynamics.

One of the purposes of this experiment was to find a positional range for the temporal overlap between pump and probe pulses on the delay stage. This will aid in future experiments on the free-space interaction that the proof-of-concept ponderomotive requires. A lower bound for time zero was approximated by first looking at the region between -50 ps and 100 ps. The rise of the sigmoidal shape can itself be split into two components: an initial slow rise and the rapid formation of the final signal. The zero-intercept (blue line in Fig. 6) of a line fit (dashed red line) to the points located at the slow-rise timepoints provides the nominal location of time zero on the delay stage (-83.1 ± 0.9 mm). Other interpretations have placed time zero at half of the sigmoidal rise^29^, though, as previously mentioned, the relationship between rise time and time zero is not well understood. We will use the half-sigmoid rise time, which corresponds to a position approximately 21 mm away from our initial time zero, as an upper bound. The error in the stage position is given by the lines (solid red lines) fit to the errors to the mean incorporated into the data. As a check, this error is compared to the error associated to the line fit itself, with the result being identical within a 95% confidence interval. These curves are drawn with the assumption that the electron pulse duration is smaller than the separation between timepoints.

**Fig. 5.**
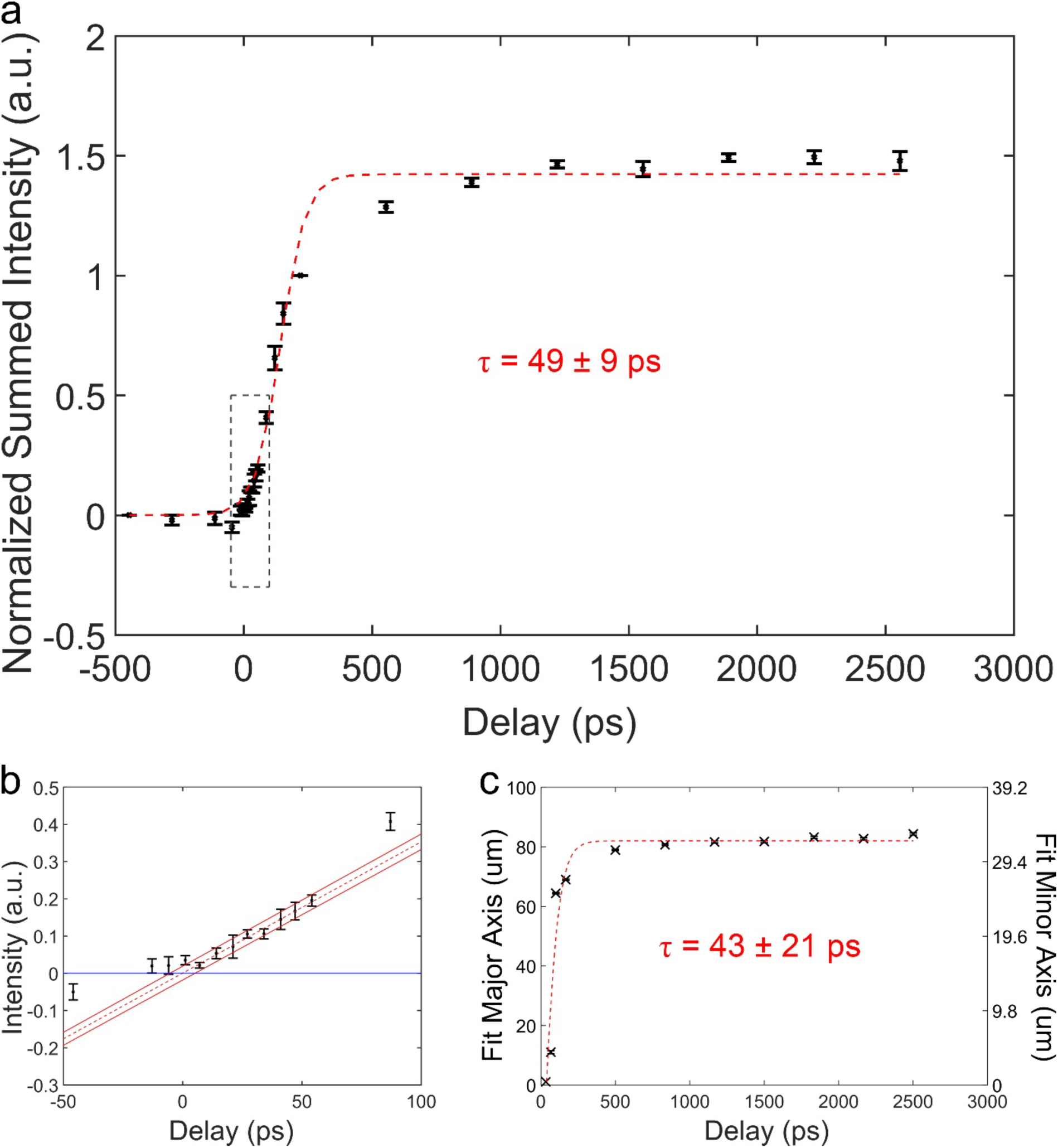
Normalized summed intensity of the difference images over delay and shape analysis. a, The photoexcitation response is plotted in discrete timepoints with error bars as standard errors to the mean. For each trial, the normalized summed intensities were normalized to two fixed points on the motorized delay stage (-446 ps and 221 ps), causing the errors at those points to be zero. A sigmoidal fit to this data with a characteristic rise time of 49 ± 9 ps is plotted as a red dashed line. An expanded view of the region bounded by the small, dashed box is provided in the bottom plot. **b**, A series of red lines are provided to demonstrate the slope of the rise in the signal, with the solid lines bounding the error in the fit. The point where the dashed red line intercepts the horizontal blue line is where time zero is approximated. **c,** The distance from the center of the annulus where the intensity, averaged over an ellipse similar to the laser spot, is at its maximum. The rise time of an empirically fit sigmoid is 43 ± 21 ps.

To reiterate, the sigmoidal fit shown in Fig. 5a is empirical. If the signal is a direct result of minority carrier photoexcitation and dynamics, one would expect a convolution of the gaussian photoexcitation pulse and the exponential decay often associated with charge carrier relaxation^27,28^. From the vendor-reported resistivities of the specimen, the expected doped charge carrier concentration is 2 × 10^19^ to 2 × 10^20^ holes *per* cm^3^. This directly translates to a minority charge carrier recombination time of 100 ns^35^. Despite the 38% increase in counts, the minority carrier concentration only increases by 10^18^ cm^-3^. There is some discussion on the charge carriers inducing a band-bending effect at the oxide-semiconductor interface which increases the SE emission, but further experiments are required to fully understand the large change in counts^36^.

Beyond the temporal properties of the signal amplitude, the shape of the signal itself also develops as a function of time. This data was calculated by first drawing an expanding ellipse from the center of the annulus. As the ellipse expands, the average intensity is recorded as a function of axis length and time (Supplementary Fig. 2b, c). For each timepoint, the signal from the expanding ellipse is then split into two portions: a rise from the center of the deficit zone and a decay at the outer edges. A plot of this peak position is provided in Fig. 5c, where we see another sigmoidal fit with a characteristic rise of 43 ± 21 ps. However, when comparing with the charge carrier diffusion length of 20-30 cm^2^/s, this value is far in excess of what could be expected for the charge carriers to travel in the same amount of time^37,38^. Indeed, taking the peak distance of 81 µm from the edge of the beam radius (30 µm) as the diffusion length and 43 ps as the time, the expected diffusion coefficient for this rapid expansion would be on the order of 6 × 10^5^ cm^2^/s. Furthermore, the signal outside of the laser spot is not generated by the pump laser pulse itself, as the fluence 81 µm away in the major axis is 5 × 10^-3^ µJ/cm^2^, far below the values required to see any signal at all^36^. The peak distance by itself is three times the size of the laser spot size, suggesting that the charge carriers rapidly spread through ballistic transport of hot carriers or ultrafast diffusion^31,39^. This is further corroborated by numerous publications at lower fluences where there was no reported expansion of the signal beyond the laser spot size^40–44^. With this, we are certain that we have captured real photoexcitation dynamics on the surface of the specimen.

### Future proof-of-concept ponderomotive phase plate experiment

Having established the location of time zero, the instrument can be reconfigured for phase plate experiments (Fig. 6)^22^. The difficulty presented here will be achieving high enough powers for a measurable effect, as the perpendicularly interacting beams have a weaker effect than in the established coaxial direction by an order of magnitude^45^. This is the purpose of the proof-of-concept SEM. As mentioned previously, the SEM provides a highly customizable platform to perform proof-of-concept experiments on the ponderomotive effect. All that is needed is to remove the specimen for a free-space interaction between the pulsed electrons and photons. Given that the interaction orientation (*i.e.* orthogonal pulsed beams) is identical, the only minor change required is to further narrow the laser beam so that the peak intensity is maximized during the interaction. This will enable the full proof-of-concept of the pulsed ponderomotive effect in the orthogonal orientation, as is suitable in the case of instruments that commonly lack space for optical components, such as TEMs. This orthogonal geometry was also chosen for the simplicity in hardware modifications, requiring passthrough windows as opposed to column extensions^21^. Moreover, the properties of the laser focus can be modified *ex situ*, enabling rapid modifications in response to changes in specimen and electron beam conditions. A secondary pulsed laser in a second orthogonal direction to the electron beam and first laser beam, but offset by some distance down the column, additionally offers a circular phase pattern without the discrete momentum jumps associated with Kapitza-Dirac style resonator phase plates^22^.

To reconfigure the SEM, a different detector is required. The NanoSEM is also equipped with a scanning transmission electron microscopy (STEM) mode, in which a focused electron beam is scanned across the specimen, and transmitted electrons are collected using a STEM detector. The STEM detector is still a diode collector, so a spatial filter, shown as a grating in Fig. 6, is also required to demonstrate changes in intensity in the interaction region. As the focused electron beam passes through the laser beam, the electrons will defocus, causing an intensity loss as more electrons strike the grating than the detector. After proof-of-concept is complete, the expectation is that the entire design can be transferred to the TEM provided that optical windows to the back focal plane are installed into the column. It should also be noted that the powers required to defocus the beam (second-order effect) in the proof-of-concept experiment are well-in-excess of the powers required to shift the phase by half a wavelength (zeroth-order effect), as explained through the ponderomotive effect equations^13,21,45^.

**Fig. 6.**
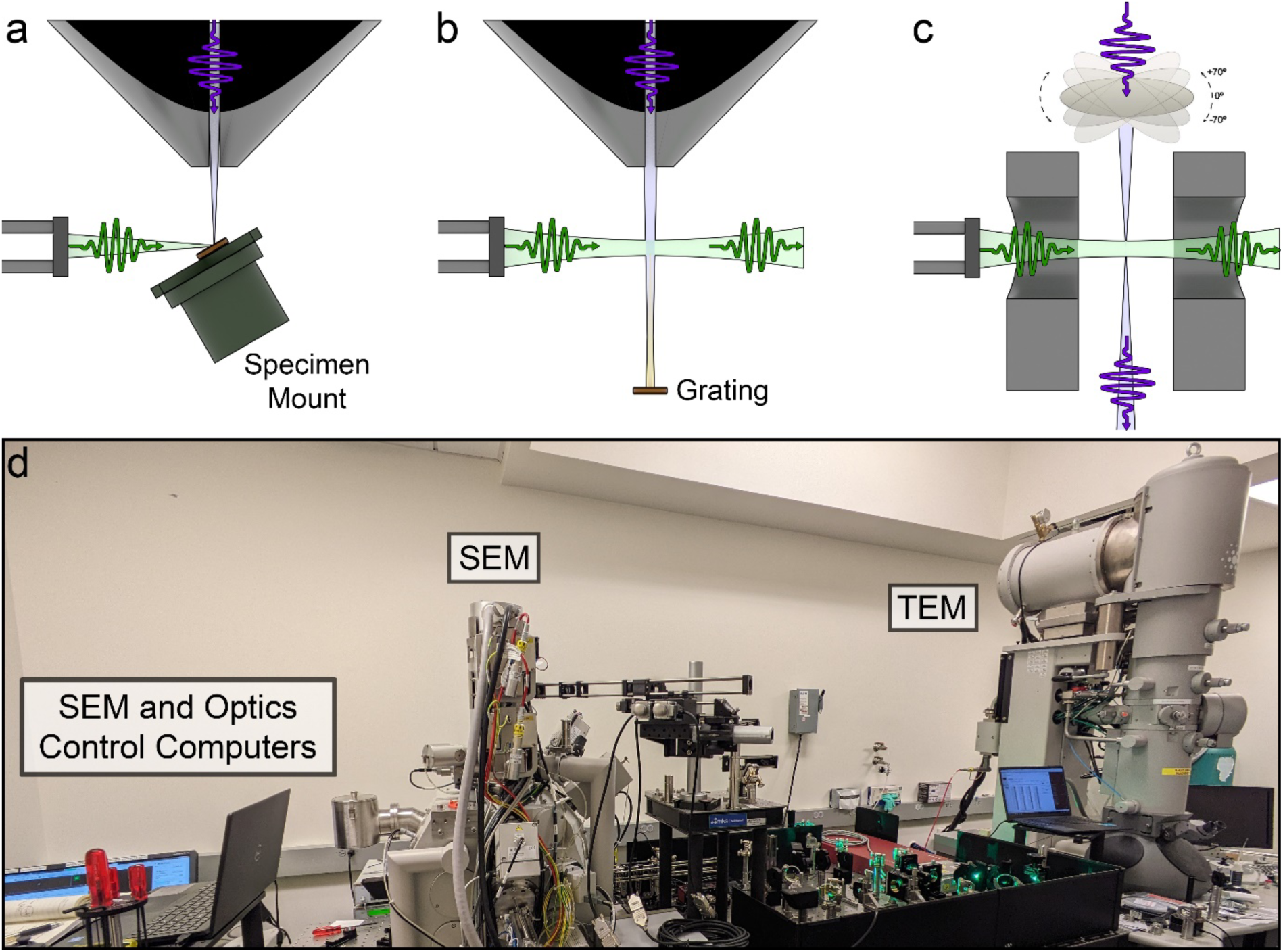
Plans for implementation of a laser-based orthogonal phase plate. **a**, A schematic of the critical time-zero experiment discussed in this paper, with the photon and electron beams focused to the same position on the specimen. **b,** The ensuing experiment planned on the very same system, with the laser now passing through the electron beam as a free-space interaction. This experiment is intended to be a demonstration of the second-order ponderomotive effect with the electron and photon pulses in an orthogonal orientation. The electron beam is focused onto a grating above a signal collection diode, with the interaction inducing an anisotropic defocusing, thus reducing the signal at the detector. **c,** The eventual implementation of the full pulsed ponderomotive phase plate on a TEM. The electron beam is initially scattered by the specimen. The photon pulses then interact with the unscattered portions of the electron pulse, shifting their phase, which then destructively interfere at the detector. **d,** The laser room in the Fitzpatrick lab with the two electron microscopes already in place. The intention is to direct the laser beam towards the TEM after proof-of-concept on the SEM is complete.

A Cryo-TEM with pulsed-laser phase plate will be similar in many ways to existing ultrafast TEMs. Like ultrafast TEMs, pulsed-laser phase plate Cryo-TEM will likely operate 1 to a few electrons *per* pulse to maximize the source brightness^46,47^. However, a critical difference exists between them: ultrafast TEMs executing light-pump, electron-probe experiments in matter have pulse repetition rates limited by the sample’s relaxation time. This typically enforces MHz-scale repetition rates and below. On the other hand, the pulsed-laser phase plate interaction with the electron beam is prompt and nondestructive. In principle, it can operate with an arbitrary pulse repetition rate, limited only by technical details of the laser, optics, and photoemission source. These details are favorable for the generation of systems with GHz repetition rates, described below.

In consideration of this repetition rate, we can first compute the required repetition rate existing photoemission nanotip sources would need to match the performance of existing continuous-wave cold field emission sources. Cold field emission (CFE) tips are the brightest source of temporally-unstructured electrons known, delivering a (reduced, independent of gun voltage) brightness of 10^8^ A/Sr/m^2^/V^48^. It has recently been demonstrated that a CFE tip brightness of 5×10^7^ A/Sr/m^2^/V enables the single-particle reconstruction of a protein complex to a resolution of 1.2 Å^49^. For pulsed TEMs operating with nanotips, the maximum reduced brightness demonstrated to date is 400 A/Sr/m^2^/V^50,51^, when operating at a repetition rate of 1 MHz. To match the brightness of continuous electron sources for cryo-EM^49^ using existing nanotip emitters, the repetition rate must be scaled up by ∼1×10^5^, or from 1 MHz to 100 GHz, which is extremely challenging. High average power, short pulse laser systems have already been commissioned in the few-GHz regime^52^ for photoemission electron accelerators with 10s to 100s of nanojoules *per* pulse; such systems could be optically split and delayed to produce >10 GHz laser systems with >10 nanojoules *per* pulse, which may be sufficient to generate a π/2 phase shift in the objective back focal plane without stringent optical focusing requirements^23^. Yet, even with 10 GHz repetition rate, this still leaves an order of magnitude in brightness between this hypothetical electron source and state of the art field emission sources.

One exciting avenue to further increase the brightness is to tune the photon’s energy to closely match the material’s work function. This can dramatically reduce the emitted electron momentum spread, particularly when the emitter itself is cryogenically cooled; energy spreads an order of magnitude below traditional field emission from metal have been demonstrated^53,54^, where here emission brightness is inversely proportional to this energy spread. To mitigate heat load on the emitter, high quantum efficiency emitters are required, particularly with prompt emission time; thin film, epitaxial alkali antimonide photocathodes are a promising candidate emitter material^55^. Yet, nanoscale emission areas with these materials have yet to be demonstrated and this is an important avenue of future research and development. Such a hypothetical source could provide the brightness of a cold field emission gun but with temporal electron control and full freedom to shape the optical phase-plate wavefront in the back focal plane of the objective lens.

In the near term, it may be possible to achieve a laser-pulsed CFE brightness sufficient for cryo-ET by simply adjusting existing tilt-collection schemes (Table 1). Assuming an optimal CFE brightness of 5×10^7^ A/Sr/m^2^/V^49^, dose-symmetric tilt schemes^5^ using fine sampling of 1°, 2°, or 3° increments would require experimental parameters like those outlined in Table 1. Under these standard conditions, acquisition times *per* tilt are on the order of tens of milliseconds.

**Table 1.**
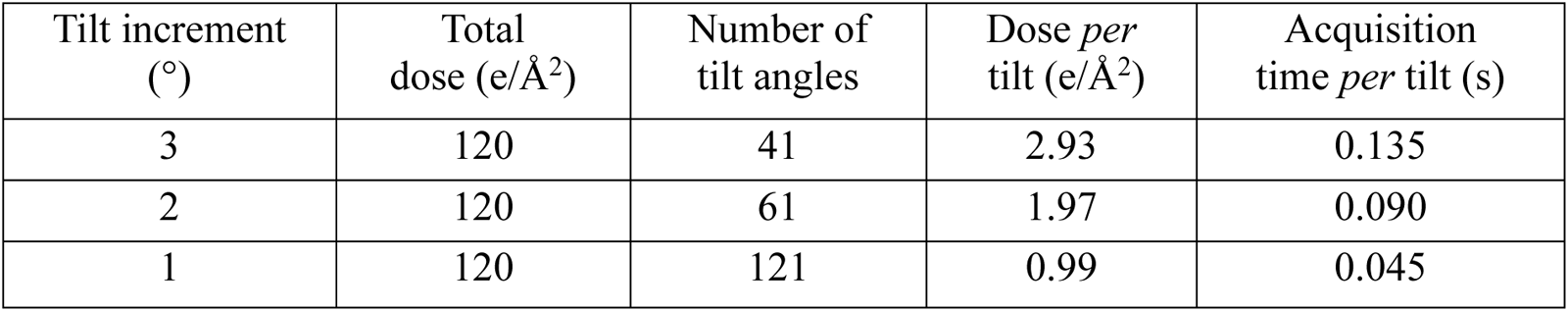
Tomogram acquisition parameters for conventional cryo-ET.

This acquisition time is remarkably fast and could be lengthened to accommodate a less bright, but equally coherent, laser-pulsed source (Table 2). If the acquisition time *per* tilt is increased by a factor of 10, a short exposure of 1.35 s is eminently practicable and would not significantly lengthen tilt-series acquisition rates^56^. For tilt increments of 3°, this reduces the required source brightness by a factor of 10 to 5×10^6^ A/Sr/m^2^/V. If the acquisition time *per* tilt is held constant at 1.35 s, the required source brightness using finer sampling of 2° or 1° tilt increments are 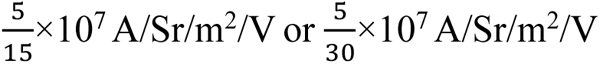, respectively, due to the lower dose *per* tilt requirements (Table 2). In principle, this would reduce the 100 GHz laser pulse rate – at most – by a factor of 30 to only a few GHz, which is within the specification limits of commercially available pulsed lasers^57^.

**Table 2.** Tomogram acquisition parameters for laser-pulsed cold-FEG cryo-ET.

| Tilt increment (°) | Total dose (e/Å<sup>2</sup>) | Number of tilt angles | Dose *per* tilt (e/Å<sup>2</sup>) | Acquisition time *per* tilt (s) | Brightness required (ASr<sup>-1</sup>m<sup>-2</sup>V<sup>-1</sup>) |
| --- | --- | --- | --- | --- | --- |
| 3 | 120 | 41 | 2.93 | 1.35 | $5.00 \times 10^6$ |
| 2 | 120 | 61 | 1.97 | 1.35 | $3.33 \times 10^6$ |
| 1 | 120 | 121 | 0.99 | 1.35 | $1.67 \times 10^6$ |

With these parameters, we have laid out a path for a pulsed laser phase plate using current-day materials and techniques. Having demonstrated an operational SUEM with a photoexcitation experiment, we now turn our attention to the proof-of-concept experiment. By demonstrating the ponderomotive effect in an orthogonal interaction, we will open the door towards realizing a π/2 phase shift in a TEM with *ex situ* control of all required laser parameters.

## Methods

### Scanning ultrafast electron microscopy

The SUEM was only modified once beyond the pump port window leading to the specimen: the optical port at the cathode region was replaced with a Torr Scientific VPZ16QBBAR UV-AR coated window to prevent multiple smaller temporal lobes from being developed in the electron signal. The UV beam is initially focused onto the cathode through the long lever arm leading to the optical port. Pulse energies of 7.5 nJ were used for a strong photoelectron signal ^50,58^. Irises are placed on this arm at the very end just after the final lens and just after the final mirror attached to the arm. The tip can then be aligned through both irises by dropping the heating current running through the tip so that the cathode is barely incandescent. A piece of paper can be placed in front of the optical port to align the UV through the same irises. We have found that this rough alignment system is capable of aligning the UV beam to within 50 microns of the tip, so a coarse motorized scan of 10-micron steps can easily cover the remaining distance, with 1 micron steps to further refine the signal. After alignment is complete, the tip can be further cooled (1 ampere heating current) to further suppress thermionic emission. The SUEM must also be tuned such that the electrons are focused through the column apertures. On the NanoSEM, this was done by modifying the initial condenser lenses (*via* service software and the default UI) such that the intensity of the photoelectrons was maximized. In most cases, spot size 1 was the most optimal condition for achieving high electron intensities. If all condenser lenses are not tuned, then there are several orders of magnitude of loss to the sides of the column and the initial electron signal will be incredibly weak. Like other groups, we found that the gradual loss of ZrO on the tip would diminish the electron signal over a 30–60-minute period^29,31,33^. A brief 5–15 minute reheat current set to factory recommended values would restore this signal for another set of images.

There are also two peculiarities with regards to data acquisition. The first is that there may be a banding effect in the actual electron signal that occurs at 15-17 Hz. In the case of a long pixel dwell time, an issue arises where there are strong bright and dark bands that inevitably increase the noise in the image, even as integration time also increases. To mitigate this, faster dwell times (≤ 1 µs) are required (Supplementary Fig. 3) to smooth out the banding effect over a large area of the image. During acquisition, the field-of-view is averaged 32-256 times with a further integration of 40 of such averaged images *per* timepoint, with the result being a high-quality image of the specimen (Supplementary Fig. 4). Additionally, there is a small amount of signal from the pump itself entering the ETD. If the initial histogram of the raw images is within the bounds of the minimum and maximum values on the detector, this pump signal is easily subtracted, though a thin filter can be placed between the light pipe and the detector itself.

Specimen preparation was minimal. The specimen (MTI p-type boron-doped Si <100> with 0.001 to 0.005 Ω·cm resistivity) was placed in baths of acetone, isopropanol, and de-ionized H_2_O in succession after 15 minutes in each. The specimen was transferred to the vacuum within 10 minutes. Similar to other papers, we found no need for a dip in hydrofluoric acid^33^. The size of the laser beam was measured with a knife-edge test using the specimen stage and a deconvolution between a gaussian and a semi-infinite wall.

### Data Analysis and Normalization

Each trial cycles back and forth through the timepoints twenty times. Each cycle contains two images of each timepoint, with each image as described above. The images pertaining to each timepoint are summed and sorted before any further image processing is done. After summation, any extraneous banding from the aforementioned banding effect is removed by subtracting the mean of all horizontal pixel lines. An additional median filter (5 px radius) was used to visually smooth the signal, though it had no appreciable effect on the signal amplitudes. At this point, the image of the specimen is clearly visible, and a quick normalization step is done to keep the values within an expected range. However, to obtain a good comparison from image to image, they must be normalized not only between 0 and 1, but also to the reference timepoint. A 100 pixel × 100 pixel fraction of the image is taken from the bottom right corner, as far away from the dynamics as possible. Each timepoint image is divided by the width of the histogram from this region to match intensities as much as possible. For each trial, the difference series is obtained, then averaged with the difference series of all other trials by following the normalization technique described in the results. As a requirement, the timepoints before time zero must have a histogram with a mean as close to zero and as narrow as possible to deem this normalization step a success. This same criterion applies to locations in the image far away from the laser spot. It is important that the difference series of each trial is acquired before averaging across trials, as a full summation across all trials will cause extraneous artifacts to appear in the final difference image (Supplementary Fig. 5). Five trials of 40 images *per* timepoint were acquired in the time range between -446 ps and 221 ps. Seven were acquired after 221 ps. In all trials, both the -446 ps and 221 ps timepoints were included.

## Acknowledgements

This work was supported by the Chan Zuckerberg Initiative, Visual Proteomics Grant under award no. 2021-234816.

## Author contributions

D.X.D., A.C.B., C.J.R.D., E.N., D.Y., J.M.M., and A.W.P.F. conceived the experiments. D.X.D., T.T., C.S., and A.W.P.F. designed and fabricated components. D.X.D. built the optical set-up and carried out the experiments and data analysis, supported by A.C.B., U.C., E.N., D.Y., J.M.M., and A.W.P.F. The manuscript was written by D.X.D., A.C.B., J.M.M., and A.W.P.F. after discussions with, and input from, all authors. J.M.M. and A.W.P.F. supervised the project.

## Competing interests

The authors declare no competing financial interests.

---

**Supplementary Fig. 1.**
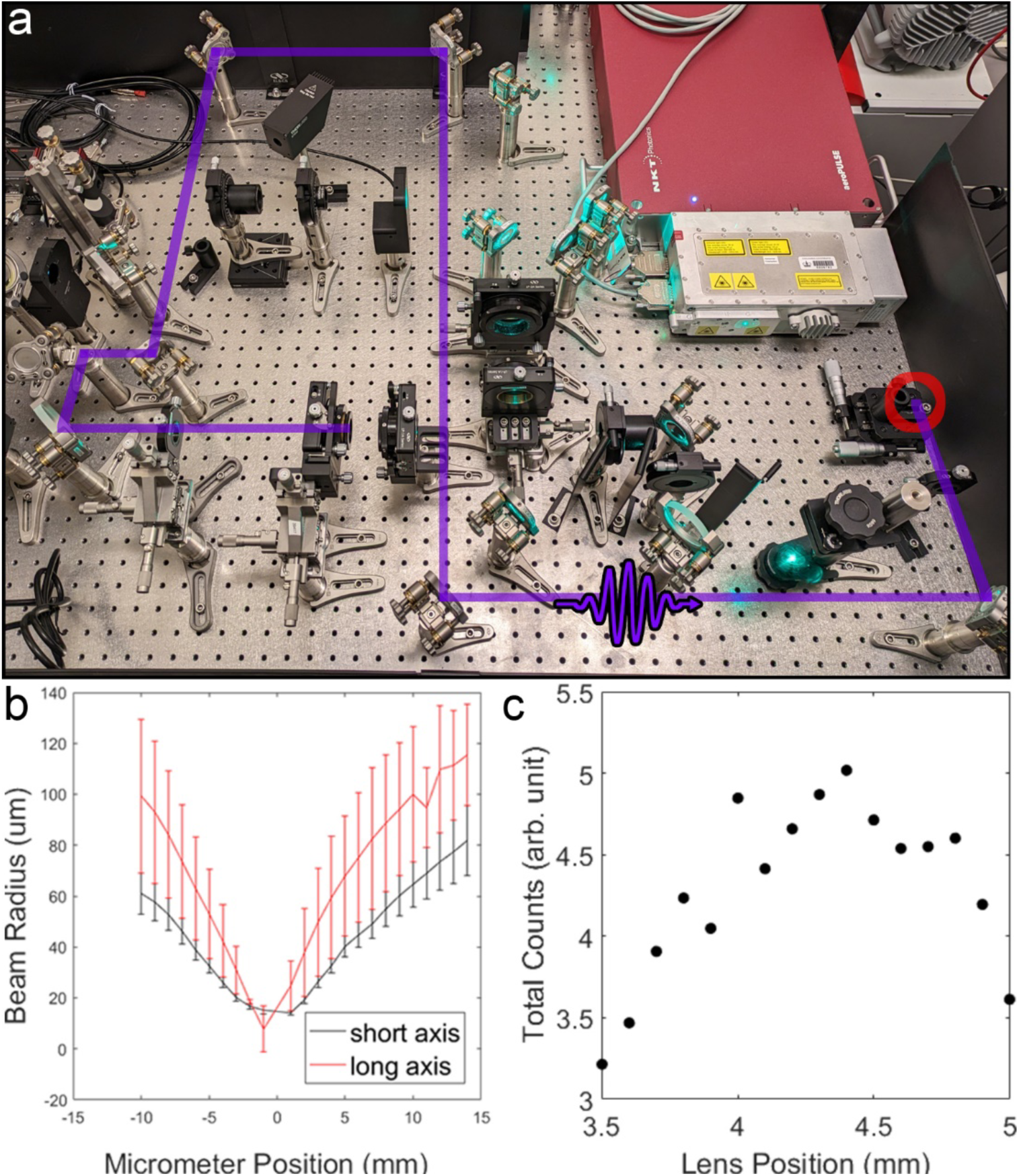
Characteristics of the UV beam and associated photoelectron emission measured as counts on a TIFF image. **a**, The path length of the UV beam was replicated on the laser table, with a lens placed in a position similar to the one found right before the optical probe port at the top of the SEM column. A beam profiler is placed at the red circle and the position is varied along the optical axis to measure the effective focal length. **b,** The size of the beam as a function of micrometer position of the beam profiler is plotted. The minimum size of the beam is 7 µm × 14 µm. **c,** The intensity of images on the STEM detector as a function of position from the final lens stage. There is a point along the stage where the intensity reaches a clear maximum, indicating the focal point of the UV beam has reached the cathode.

**Supplementary Fig. 2.**
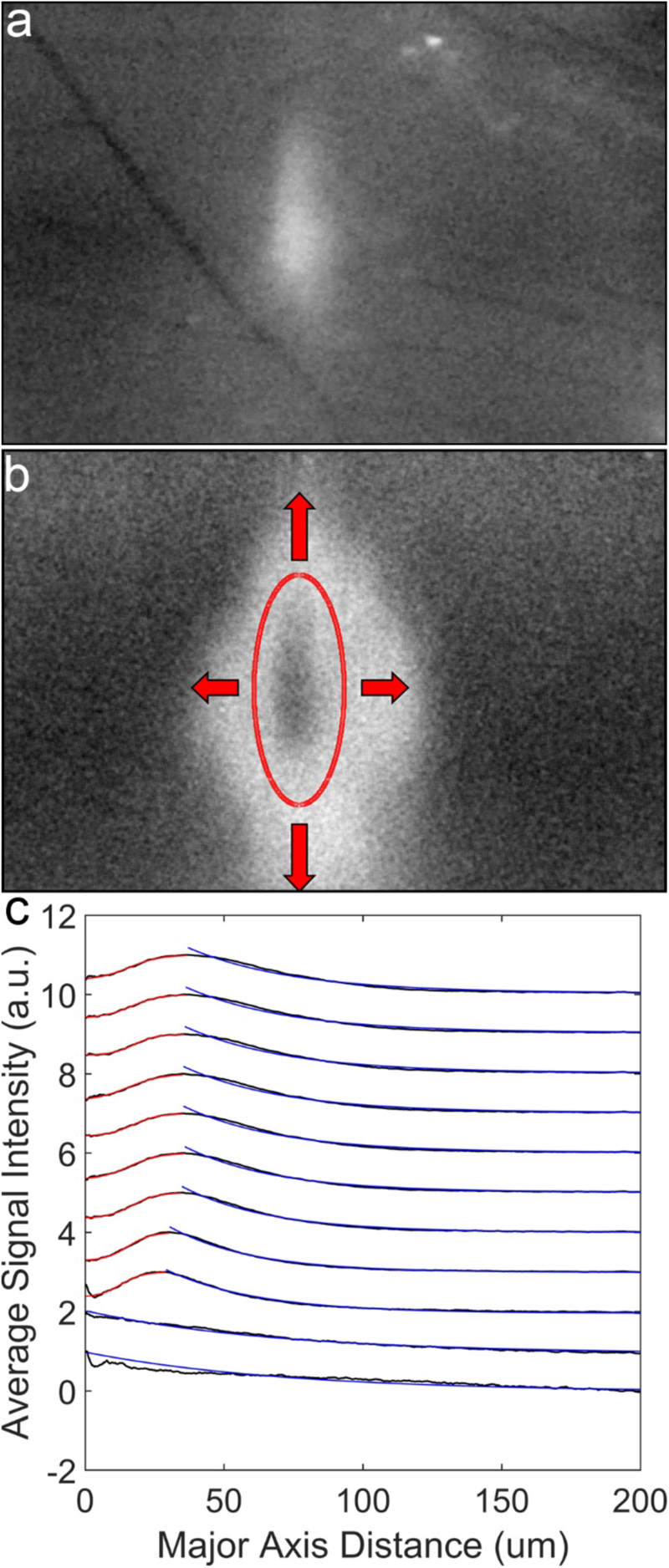
Measuring the rise/decay of signal for each timepoint. **a**, A reference image of the specimen well before time zero. **b,** An ellipse with a major/minor axis ratio equivalent with the major/minor beam waist ratio of the pump beam is drawn centered on the signal. The ellipse is then expanded until an arbitrary distance of 200 microns in the major axis direction is reached. For each ellipse, the signal is integrated and then normalized to the total number of pixels. **c,** The results for each timepoint beginning from 87 ps and onwards, artificially separated to more easily visualize differences in the signal. The rise of the signal from the center is fit to a sigmoid and the decay is fit to an exponent. The characteristic lengths are shown in Fig. 6.

**Supplementary Fig. 3.**
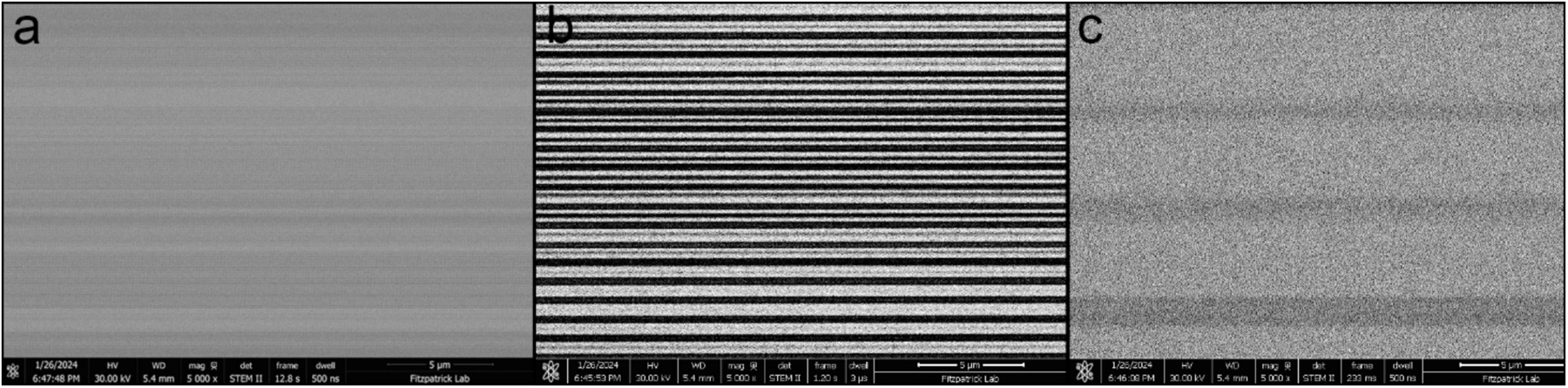
Demonstrating the acquisition conditions necessary for coherent signal after image processing. **a**, A high-resolution image at 500 ns of dwell time showing a banding effect. The frequency of this banding is approximately in the 15-17 Hz range, with no clear reason for its appearance. **b,** If higher dwell times are used, as in the case of this 3 µs dwell time image, the additional noise from the off-band regions essentially becomes a loss in signal. High enough dwell times will cause incoherent noise to cover the image, regardless of integration time. **c,** To mitigate this effect, shorter dwell times are used to smooth the bands into large areas across the image. This causes more coherent accumulation of signal and any additional banding effects afterwards can be removed by subtracting the means across the horizontal direction.

**Supplementary Fig. 4.**
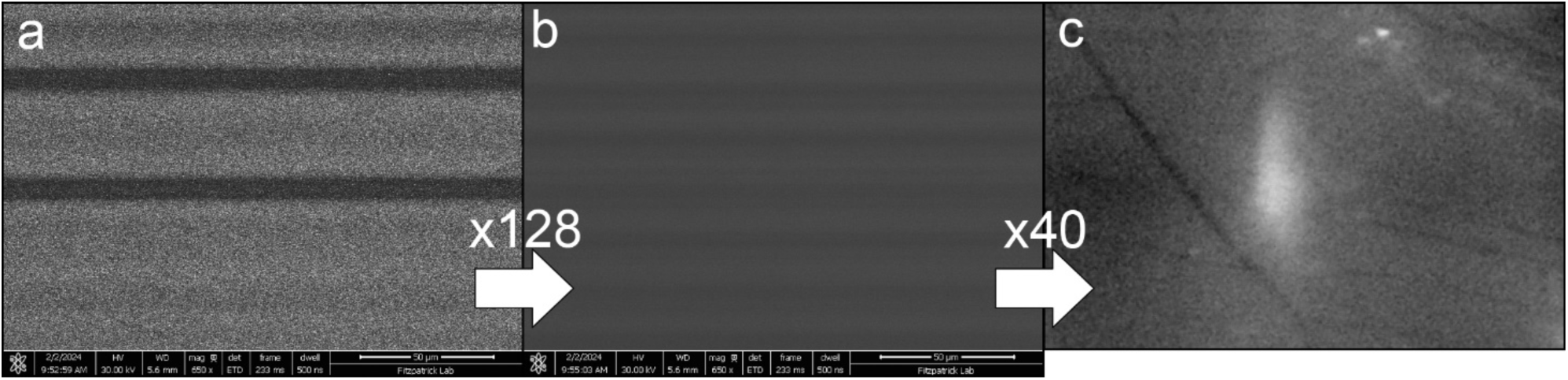
The number of integration steps required for a full image in a single trial. **a**, A purely raw image acquired at 500 ns of dwell time. This raw image is averaged 128 times during image acquisition by the SEM software to form **b.** 40 of these images are collected *per* trial, resulting in a total integration time of 1000 seconds. **c,** There is a significant banding effect in the final image, so the mean of each horizontal line is subtracted for the full result.

**Supplementary Fig. 5.**
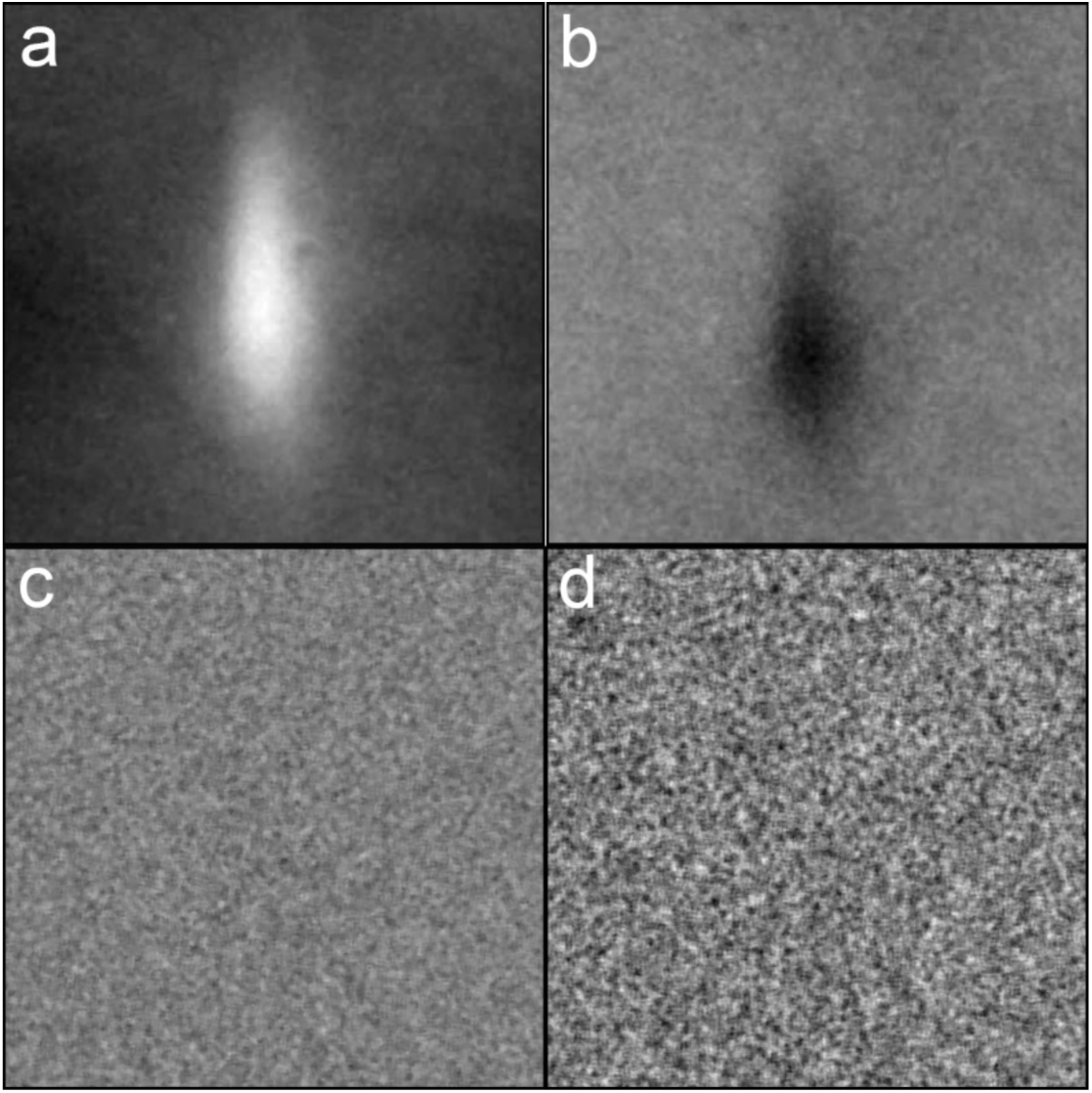
Order of operations in image processing is critical. **a**, The sum of all of the images in a single timepoint across all trials. The object in focus is the damage region caused by the pump beam. **b,** When subtracting the images from the reference image as a single set, a dark signal remains. This is directly due to the normalization technique, which can lead to remaining spots on the difference images from small differences between each trial. **c,** The difference image in a single trial at the same timepoint, showing no signal in the exact same region. Calculating the difference images on a trial-by-trial basis is critical for ensuring no incoherent ultrafast signals leak through in the final analysis. **d,** The averaged difference image from every single trial in the timepoint. Here we see that, despite using the same number of images as (b), non-ultrafast signals are accurately removed.

## Notes

### Competing Interest Statement

The authors have declared no competing interest.

